# The effect of an ultrasound-activated electrospun piezoelectric hydrogel scaffold on post-traumatic brain injury motor function in *Drosophila melanogaster*

**DOI:** 10.64898/2026.09.20.752928

**Authors:** Mihir Nimkar, Ronav Gopal

## Abstract

Traumatic brain injury (TBI) is a leading cause of long-term neurological disability, affecting 50-60 million people annually. Cascading secondary injury mechanisms, including oxidative stress and neuroinflammation, impair motor and cognitive function, while current therapies often manage symptoms rather than restore lost neurological function. This study investigated an ultrasound-activated piezoelectric hydrogel scaffold composed of barium titanate nanoparticles (BTNPs), sodium alginate (SA), and polyethylene oxide (PEO). The hypothesis was that TBI flies receiving the scaffold and ultrasound would display a higher climbing assay pass rate than TBI flies receiving no treatment or BTNPs alone. TBI was induced in *Drosophila melanogaster* with a high-impact trauma (HIT) apparatus. Treatments were applied by opening and resealing the fly head cuticle, followed by ultrasound stimulation. Motor recovery was quantified with a climbing assay across four groups (N=10 flies per group), which measured the percentage of flies that could cross a 5 cm mark within 30 seconds. Assay validity was supported by a statistically significant difference between flies without TBI and those with TBI using Mann-Whitney U testing (p=0.0035). Although TBI treatment groups did not differ significantly (p>0.05), median pass rates increased from injury control (35%) to BTNPs (45%) to scaffold (50%). However, high inter-trial variation, a 45.2% procedural mortality rate, and small sample size limited statistical power. These results suggest that piezoelectric electrospun hydrogel scaffolds may be a pathway for safer and more biocompatible restoration after TBI, but provide insufficient evidence to conclude that the scaffold significantly improved motor recovery.

## Introduction

Traumatic brain injury (TBI) is a leading cause of long-term neurological disability and death worldwide, affecting approximately 50 to 60 million individuals each year and generating an estimated $400 billion in global economic losses annually (Zhou et al., 2025). Common causes include falls, motor vehicle accidents, violence, and sports-related injuries. Despite its prevalence, few restorative therapies effectively address long-term neural damage after the acute phase has passed. As highlighted in Figure 1, the pathology of TBI is typically divided into primary and secondary injuries. Primary injuries are caused by direct mechanical forces, including white matter damage, tissue shearing, contusions, skull fractures, concussions, and tissue compression. While primary injury is partly managed through surgical intervention and acute pharmaceuticals, these approaches do not address long-term neural repair or tissue regeneration (National Institute of Neurological Disorders and Stroke, 2025). Secondary injuries are caused by molecular cascades triggered by initial impact and include neuroinflammation, cognitive and motor dysfunction, and accelerated neurodegeneration through oxidative stress (Katzenberger et al., 2013). Oxidative stress occurs when reactive oxygen species (ROS), unstable molecules with unpaired electrons, bypass antioxidant defenses to incite chemical reactions that damage cells. Cell death contributes significantly to secondary injury progression; therefore, enhancing the regenerative potential of neural stem cells (NSCs), which are capable of differentiating into functional neurons and glial cells, is an important target for improving recovery following TBI.

**Figure 1.**
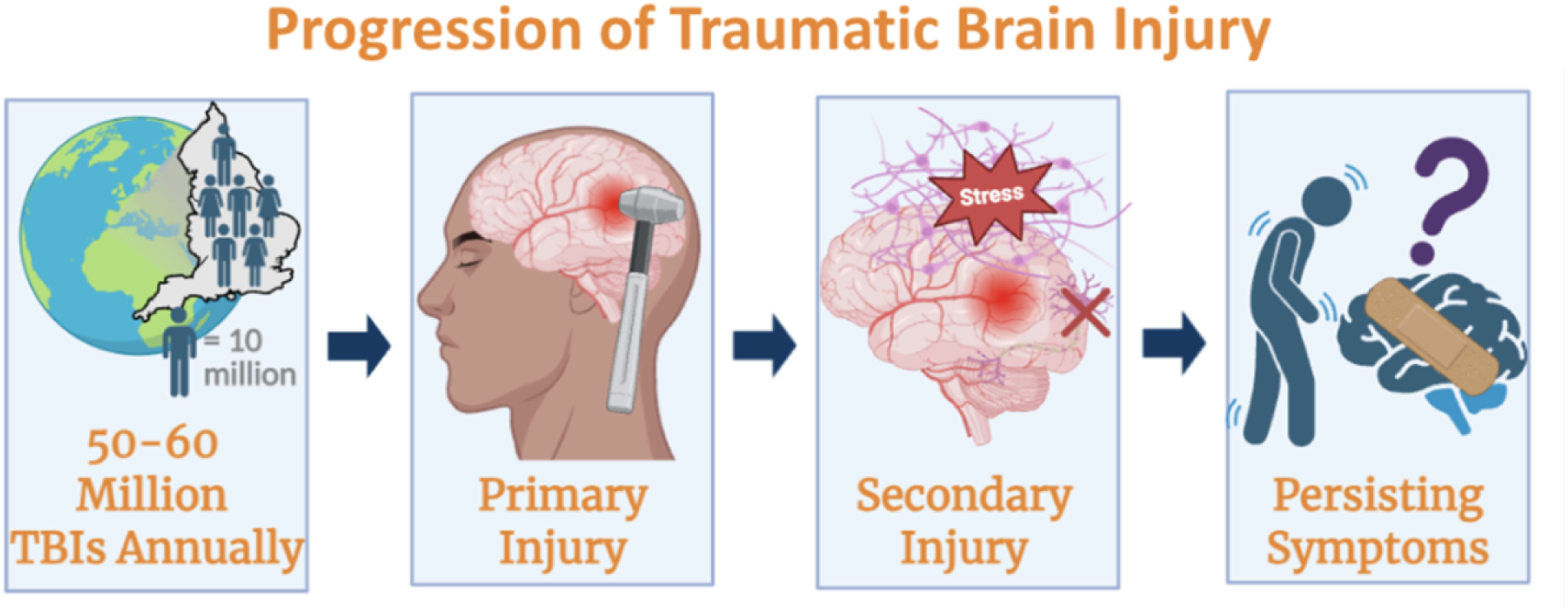
Diagram showing the progression from primary mechanical injury to secondary injury mechanisms after TBI. Primary injury includes tissue shearing, contusions, compression, and white matter damage, while secondary injury includes oxidative stress, neuroinflammation, cell death, and impaired neural signaling. Created with BioRender.

Current treatments demonstrate mixed clinical outcomes across patient populations and often focus on limiting acute damage rather than restoring long-term neural function. Cellular therapy involving stem cell transplantation has shown limited improvement in speech, cognition, or memory relative to controls in recent clinical trials, partly because NSCs differentiate slowly under physiological conditions and cannot efficiently reconstruct neural networks on their own (Zhou et al., 2025). Intracranial implantation and cell injections of cellular therapy are also invasive, expensive, and carry surgical risk. So, although these natural regeneration mechanisms appear to be crucial, supplementing them with other methods seems to be necessary for any effects. Growth-factor based therapy faces a separate limitation because growth factors may diffuse away from the injury site, resulting in reduced localization and decreased ability to direct targeted cell differentiation (Wang et al., 2025). Pharmaceutical therapies have also had high failure rates across multiple studies of neuroprotective drugs. Pharmaceutical therapies also often have a narrow treatment window limited to only hours after the injury and largely focus on the acute phase of TBI such as reducing swelling or inflammation rather than long-term cognitive and neurological deficits (Zhou et al., 2025).

Another emerging approach relevant to this study is neuromodulation to potentially ameliorate TBI symptoms. Although neuromodulation has primarily been used in TBI-induced comas, certain methods of electrical stimulation have been used to alleviate motor issues in TBI (Zhou et al., 2025). Specifically, piezoelectric devices harvest ultrasound energy for brain stimulation, leading to deeper and more targeted brain activation than electrical stimulation (Hou et al., 2024). Piezoelectric nanoparticles can generate electrical signals when stimulated by ultrasound and have been applied to promote regeneration in NSCs (Wang et al., 2025). However, this promotion still relies on the transplantation of stem cells and can be invasive or costly, showing the need for a more direct method that supports natural healing rather than introducing new cells while also being more effective. This study tested whether embedding barium titanate nanoparticles (BTNPs) within an electrospun SA/PEO hydrogel scaffold could provide a more structured and biocompatible format for ultrasound-activated stimulation after TBI. By using electrical stimulation with external ultrasound, this study addressed limitations of freely delivered nanoparticles by applying piezoelectric stimulation in a scaffold-based format.

A recent study investigated the use of ultrasound-activated piezoelectric stimulation in improving NSC-based therapies for treating TBI. Specifically, barium titanate (BaTiO_3_) nanoparticles with a high piezoelectric coefficient and strong biocompatibility have shown promise in performing piezoelectric stimulation for neural differentiation and nerve regeneration. However, BTNPs can be engulfed by brain cells to generate ROS, which can result in cellular death and led to the piezoelectric nanoparticle therapeutic having no effect on TBI in the study. Thus, the study emphasizes the need to enhance piezoelectric stimulation-based NSC therapy by preventing ROS production. Instead of a hydrogel, however, the study used a structural material made of graphene to reduce how much of the nanoparticles were taken up by cells through the rigid structure that is harder to engulf. The study combined the structural material with the barium titanate piezoelectric nanostickers for TBI in mice and found that the nanostickers were effective, providing precedent for generally using piezoelectric nanoparticles, as well as specifically using barium titanate, when a structural component is present that can prevent cells engulfing nanoparticles and ensure biocompatibility (Wang et al., 2025).

Another piece of current research into neural therapeutics is the regenerative effects of tissue engineering for neural injuries. Specifically, neural tissue engineering can provide healing to damaged neurons and neural tissues by using various stem cells and biomaterials, which can mimic the natural extracellular environment (Fang et al., 2025). The electrospinning of hydrogel-based materials has specifically shown promise in this task. Hydrogels are injectable, biocompatible, and do not induce swelling, making them well suited for being in a physiological environment such as the brain. Hydrogels also often exert control over the neural tissue of the host to support the growth, differentiation, and replication of NSCs (De et al., 2022). Hydrogel-based scaffolds have had some precedence in promoting neural regeneration and reducing tissue damage amidst stimulated inflammatory and oxidative stress responses (Du et al., 2023).

The potential for hydrogels to reduce oxidative stress, which is caused by ROS, is important because ROS are commonly produced at biomaterial implantation sites, leading to cytotoxic effects that can cause failure of cellular therapies or even worsen TBI. A recent study found that certain hydrogels formed a ROS-protective composite (Canic et al., 2025). Hydrogel-based scaffolds can additionally provide structural support and provide targeted delivery to promote neural therapy while mitigating secondary injury. The structural support is especially important so that the barium titanate nanoparticles aren’t as easily engulfed by cells to cause ROS and cell damage as seen in some cases. Hydrogels have also been used to support other therapy methods, like when collagen-chitosan hydrogels were able to support cellular therapy that used bone marrow stem cells, increasing neurological function after implanted into a TBI lesion site compared to bone marrow cells alone by allowing for more of the stem cells to be retained and used (Du et al., 2023).

In this study, combining the piezoelectric and hydrogel components was used to test whether a structured scaffold could help reduce ROS-related concerns while also enhancing the electrical and regenerative properties of the scaffold. Specifically, sodium alginate biomaterials with piezoelectric components were coupled to form a pro-regenerative mesh. The goal of the mesh was to mimic the extracellular matrix (ECM), a three-dimensional network of proteins and surrounding cells that regulates cell behavior and provides tissue repair and regeneration. Electrospinning was used to combine the piezoelectric and hydrogel components to test whether this structure could improve the effectiveness of the treatment. Electrospun nanofiber scaffolds are often used to support materials like hydrogels, with electrical stimulation as a prospect to be combined with the nanofibers. While hydrogel scaffolds have seen some success in supporting growth factor or other biomaterials, their potential to integrate with piezoelectric materials has not been explored as extensively. Additionally, electrospinning was used to possibly increase contact area as compared to applying the materials alone, which it has been used for in previous studies (Kornev et al., 2018). If contact area was increased for more coverage, any potential electrical and regenerative properties of the scaffold may have been enhanced.

Finally, one of the best TBI models to examine the efficacy of treatments is *Drosophila melanogaster*. *Drosophila melanogaster* was chosen as a promising model for TBI because it has brain regions that are mostly homologous to human brains, with diverse neurons and similar glial cell types. TBI can be modeled in *Drosophila melanogaster* through the use of high-impact mechanical shock in the high-impact trauma (HIT) model. The HIT model involves flies in a vial attached to a strong spring that is held in place by a wooden board. An impact mat is located under the vial, and delivering high-impact trauma involves lifting the spring at a 90 degree angle and releasing, sending a shockwave through the flies to model a TBI in flies. In the study outlining the HIT model, neurological changes after the initial impact were observed to cascade into downstream secondary effects such as oxidative stress and neuroinflammation, ultimately driving observations of behavioral impairments such as decreased motor ability and some cognitive decline. Specifically, Katzenberger et al. (2013) utilized a climbing assay to assess the motor ability of the flies and found a decline in climbing ability, along with reduced lifespan, cell death, and accelerated aging, among flies induced with TBI.

Due to the similarity of conserved responses and motor symptoms in the *Drosophila melanogaster* brain, evaluating the treatment in flies was used to provide a model that can show possible implications for humans. Furthermore, while this protocol does not target only the brain, it does mirror the diversity of human TBI cases. In real life, anything from a car crash to a sports injury can cause TBI, with a vast range of forces from different directions that makes every TBI case unique. The HIT model reflects this diversity, providing a robust system that has been cited as an effective model throughout other studies (Katzenberger et al., 2013). The climbing assay used in the HIT model was also used in this study due to the many cases of use in other studies and the effectiveness of the assay for measuring motor impairment. The assay utilized negative geotaxis, the natural inclination of *Drosophila melanogaster* to climb up in a vial against gravity. This natural inclination of flies meant that an average normal fly in a climbing assay should have been able to cross a line 5 centimeters above the ground in 30 seconds easily, which is why failure to do so was used in this study to quantify motor impairment (Dollinger et al., 2017).

In order to apply the electrospun piezoelectric hydrogel scaffold to the *Drosophila* melanogaster head region, the cuticle-opening procedure was selected instead of ingestion or larval injection. While ingesting the scaffold is not supported extensively in current literature, methods to inject materials into *Drosophila melanogaster* larvae have been developed in one study. However, the study details a protocol to inject dyes into larvae with no mention of reaching the brain of the flies (Soltani et al., 2024). Because similar studies also do not use electrospun scaffolds or target the brain, using an injection protocol would have been unnecessarily costly and provided no methods to verify effective delivery of the scaffold. To mitigate the risk of not being able to verify scaffold delivery, opening the cuticles of *Drosophila melanogaster* to apply the scaffold and sealing the opening with UV glue was the method employed in this study. Although this approach allowed direct material placement, it also introduced substantial procedural stress and contributed to mortality.

The hypothesis was that an ultrasound-stimulated electrospun piezoelectric hydrogel scaffold composed of sodium alginate (SA), polyethylene oxide (PEO), and barium titanate nanoparticles (BTNPs) would improve motor function in *Drosophila melanogaster* after TBI, as measured by enhanced negative geotaxis behavior in a climbing assay. After TBI, flies receiving the scaffold and ultrasound were expected to show a higher negative geotaxis pass rate than TBI flies receiving sham treatment and TBI flies receiving BTNPs alone. This study evaluated behavioral motor recovery rather than directly measuring oxidative stress or neural regeneration. The hypothesis was supported by ultrasound stimulated piezoelectric materials that have led to some improvements in climbing ability compared to controls (Wang et al., 2025). Furthermore, there is precedence towards the idea of hydrogels reducing oxidative stress, thereby possibly improving neural regeneration (Du et al., 2023). Electrospinning has also shown promise towards healing damaged neurons and neural tissues by mimicking the natural extracellular environment, demonstrating promise towards neural regenerative capabilities for TBI (Fang et al., 2025).

This study is novel in several respects. First, it may be the first study to integrate piezoelectric nanoparticles with a hydrogel for use in the brain. Current research usually focuses on one of the other, though combining them shows promise for enhancing effectiveness. Second, the study advances the potential application of an electrospun scaffold to the brain. Current research focuses on other biomaterials that do not use fibers or combine sodium alginate with piezoelectric materials, with most of the research in the brain focusing on only one of the components. Particularly nanofibers and especially ones with these components have mainly seen use in either cell models, the spinal cord, or peripheral nerves rather than the brain. Third, research has not been done on scaffolds particularly for *Drosophila melanogaster*, suggesting new research methods to test similar remedies could be explored in simpler models like flies as compared to models like humans or rats that can be harder to work with due to their greater complexity.

The results of this study add to the field by garnering interest on electrospinning hydrogels for oxidative stress protection and cell survival neurite growth, while serving as a possible stepping stone in understanding for a therapy targeted towards TBI survivors that may be able to be adapted for less invasive delivery or become less costly. Regardless of outcome, this study provides insight into the mechanisms of piezoelectric hydrogel scaffolds, probing further research into why the combination of methods led to the result it did and how the methods work individually.

The proposed research is significant since TBI lacks disease-modifying therapies and survivors often face persistent deficits driven by secondary injury problems. Patients who cannot afford invasive treatments, complex medications, or stem cell transplants could potentially have their struggles and symptoms alleviated with a scaffold treatment that adds a new treatment into a field with few restorative therapies that are clinically used. Understanding how a complex scaffold interacts with TBI can give insight into potential uses for bioelectronics in neural therapeutics. The insight gained can not only potentially be implemented into possible direct therapeutics, but also as a method to enhance stem cell therapy, which can both advance the field greatly. Finally, future research can even expand on the results of this study to focus on other neurodegenerative diseases that cause cell death rather than TBI.

## Results

### Scaffold Characterization

Several scaffold combinations were tested before a final electrospun mesh was produced. The first trials used marine collagen peptides dissolved in PBS and ethanol and was electrospun for three attempts. However, the solution was unable to form stable fibers in a mesh and instead produced a spray-like pattern with rapid evaporation shown in Figure 2, so the formulation was deemed not sufficient for scaffold formation. Collagen was thus not used for the hydrogel component. A PEO control scaffold in water was electrospun at 18 kV, 0.5 mL/hr, and a 10 cm tip-to-collector distance, and scanning electron microscope (SEM) imaging depicted a more ideal and continuous fiber structure, suggesting that PEO improved chain entanglement and supported electrospinning. Thus, sodium alginate and PEO in water was electrospun since sodium alginate alone is difficult to spin.

**Figure 2.**
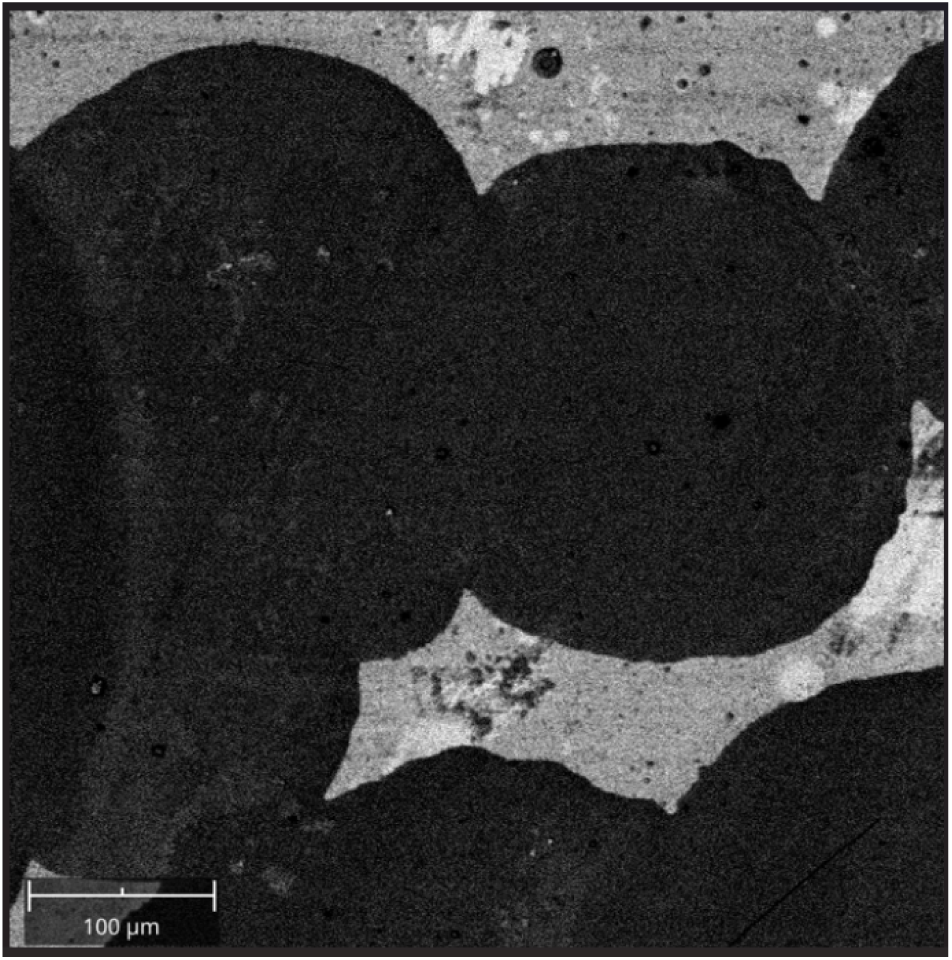
Early scaffold trials using marine collagen peptides in PBS and ethanol produced a spray-like pattern rather than stable electrospun fibers. Because this formulation did not form a usable mesh, collagen was removed from the final scaffold formulation.

Early SA/PEO scaffolds showed extensive beading, which is clumping caused by uneven material distribution during fiber formation, which forms circles called beads. Early SA/PEO scaffolds used the following electrospinning parameters: 15 cm collector distance, 0.5 mL/hr flow rate, and approximately 15-20 kV. The following scaffolds were then examined through the scanning electron microscope, which showed that clustering and beading were still visible.

Through fine-tuning electrospinning parameters, the final optimized SA/PEO/BTNP scaffold is shown in Figure 3. SEM imaging of the final SA/PEO/BTNP mesh showed reduced beading compared to the earlier SA/PEO attempts, meaning the scaffold formed a more connected electrospun structure. Although mild beading was still present, the final scaffold was considered sufficient for continued testing because it successfully formed a fibrous piezoelectric mesh and was then crosslinked with CaCl₂ and stimulated through ultrasound for therapeutic effect after traumatic brain injury. While previous meshes had smaller pores of around 4000 nm, with one having around 4600 nm pores, the final synthesized mesh had pores of around 7000 nm, which were much larger and indicated reduced beading. For the final scaffold, the fiber diameter also seemed to be approximately normally distributed as shown in Figure 3, with an average fiber diameter of around 200 nm, suggesting a nanoscale mesh well suited for biocompatibility and maximized surface area for therapeutic applications.

**Figure 3.**
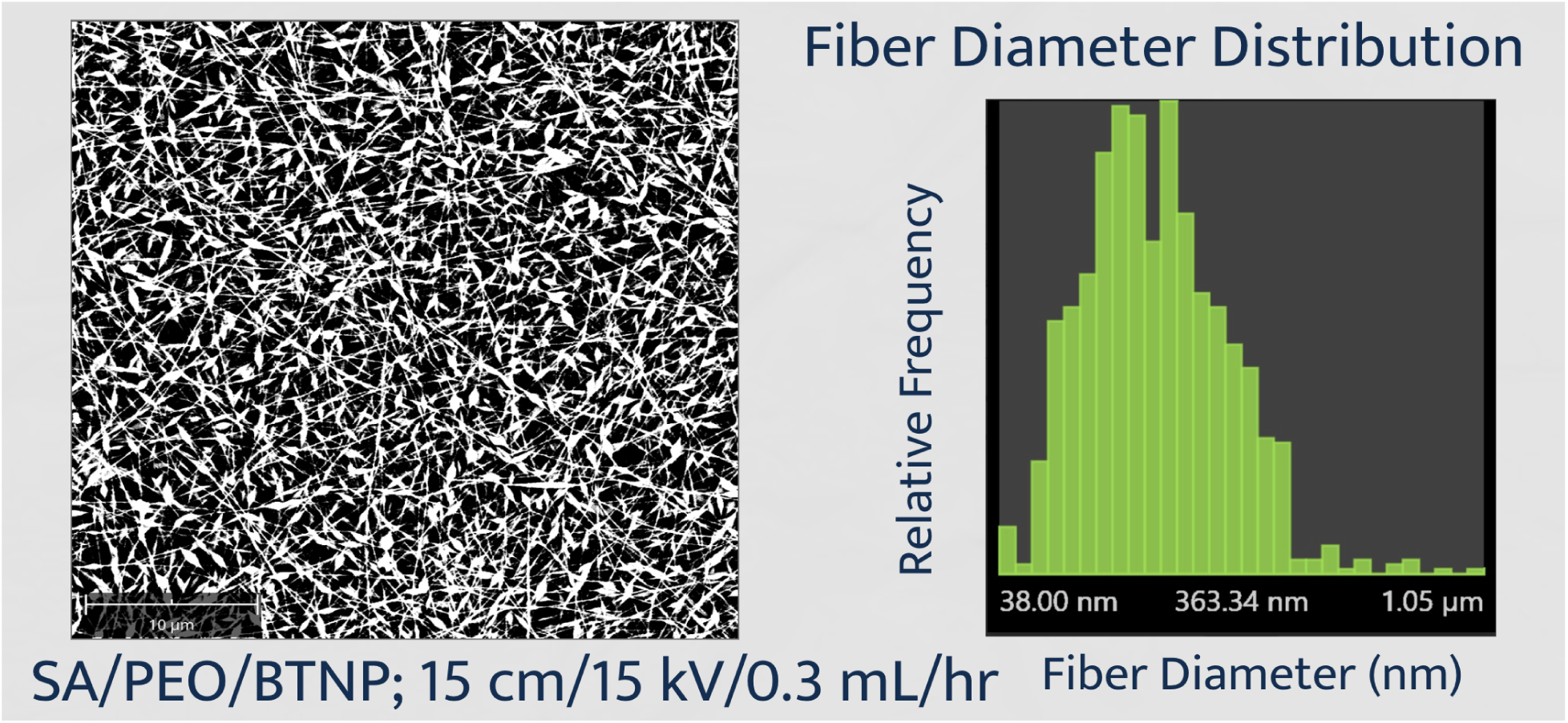
SEM characterization of the final SA/PEO/BTNP scaffold showed a connected fibrous mesh with reduced beading compared with earlier trials. The scaffold had pores of approximately 7000 nm and an average fiber diameter of approximately 200 nm, indicating a nanoscale structure suitable for treatment testing.

### Impact of Scaffold on Motor Recovery

Raw climbing assay pass percentages for all 10 trials in each of the four groups are presented in Table 1.

**Table 1.**
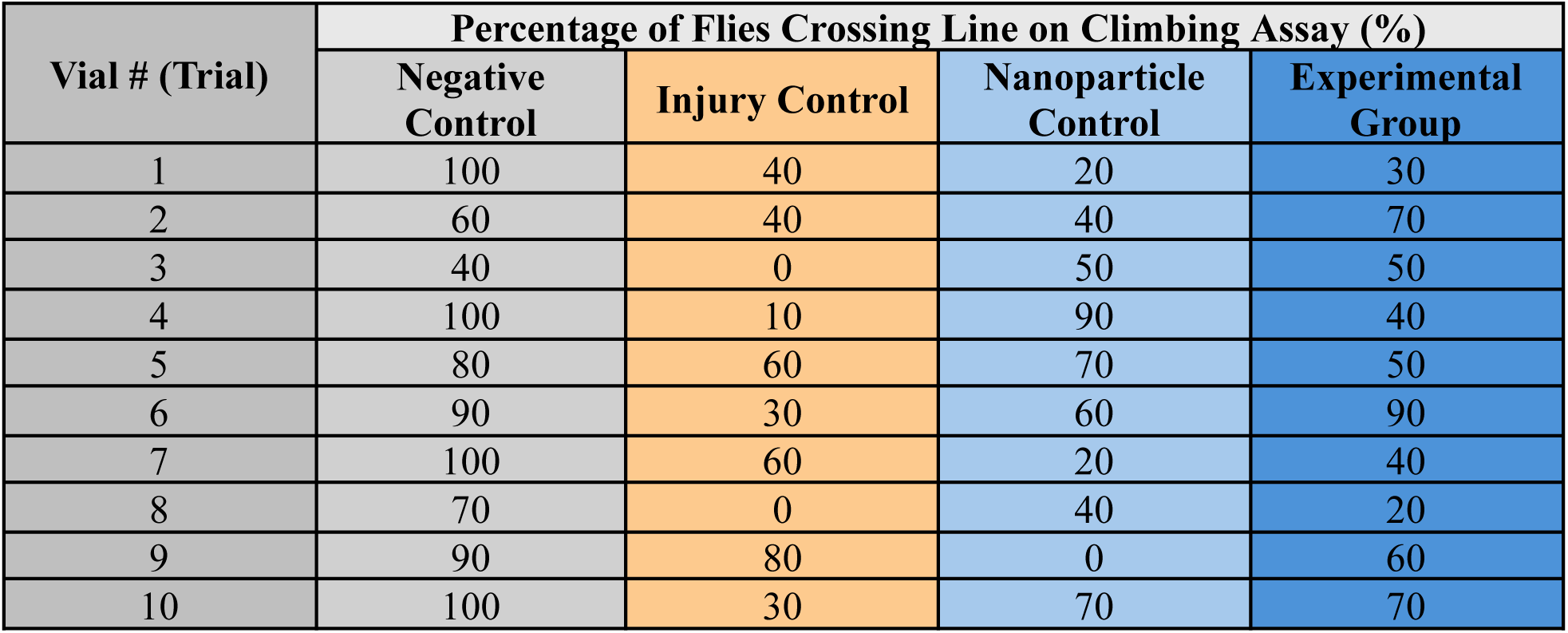
Percentage of flies successfully crossing the 5 cm mark across the four groups.

| Vial # (Trial) | Percentage of Flies Crossing Line on Climbing Assay (%) |  |  |  |
| --- | --- | --- | --- | --- |
|  | Negative Control | Injury Control | Nanoparticle Control | Experimental Group |
| 1 | 100 | 40 | 20 | 30 |
| 2 | 60 | 40 | 40 | 70 |
| 3 | 40 | 0 | 50 | 50 |
| 4 | 100 | 10 | 90 | 40 |
| 5 | 80 | 60 | 70 | 50 |
| 6 | 90 | 30 | 60 | 90 |
| 7 | 100 | 60 | 20 | 40 |
| 8 | 70 | 0 | 40 | 20 |
| 9 | 90 | 80 | 0 | 60 |
| 10 | 100 | 30 | 70 | 70 |

Prior to evaluating treatment effects, the high-impact trauma apparatus was evaluated to see if the apparatus successfully induced motor impairment and that the climbing assay was sensitive enough to detect this difference. A Mann-Whitney U test comparing the negative control to the injury control yielded p=0.0035, indicating a statistically significant difference. Similarly, conducting a Mann-Whitney U Test to compare the negative and nanoparticle controls (p=0.0060) and the negative control and experimental group (p=0.0014) yielded significant differences. As highlighted in Figure 4, interquartile ranges were 30% for the negative control, 45% for the injury control, 47.5% for the nanoparticle control, and 30% for the experimental group, indicating that variation was highest in the two intermediate treatment groups. The large IQR of 45% in the injury control group relative to the negative control IQR of 30% indicates that TBI introduced substantial within-group variability in motor performance, reflecting the severity of injury across individuals exposed to the same high-impact trauma protocol.

**Figure 4.**
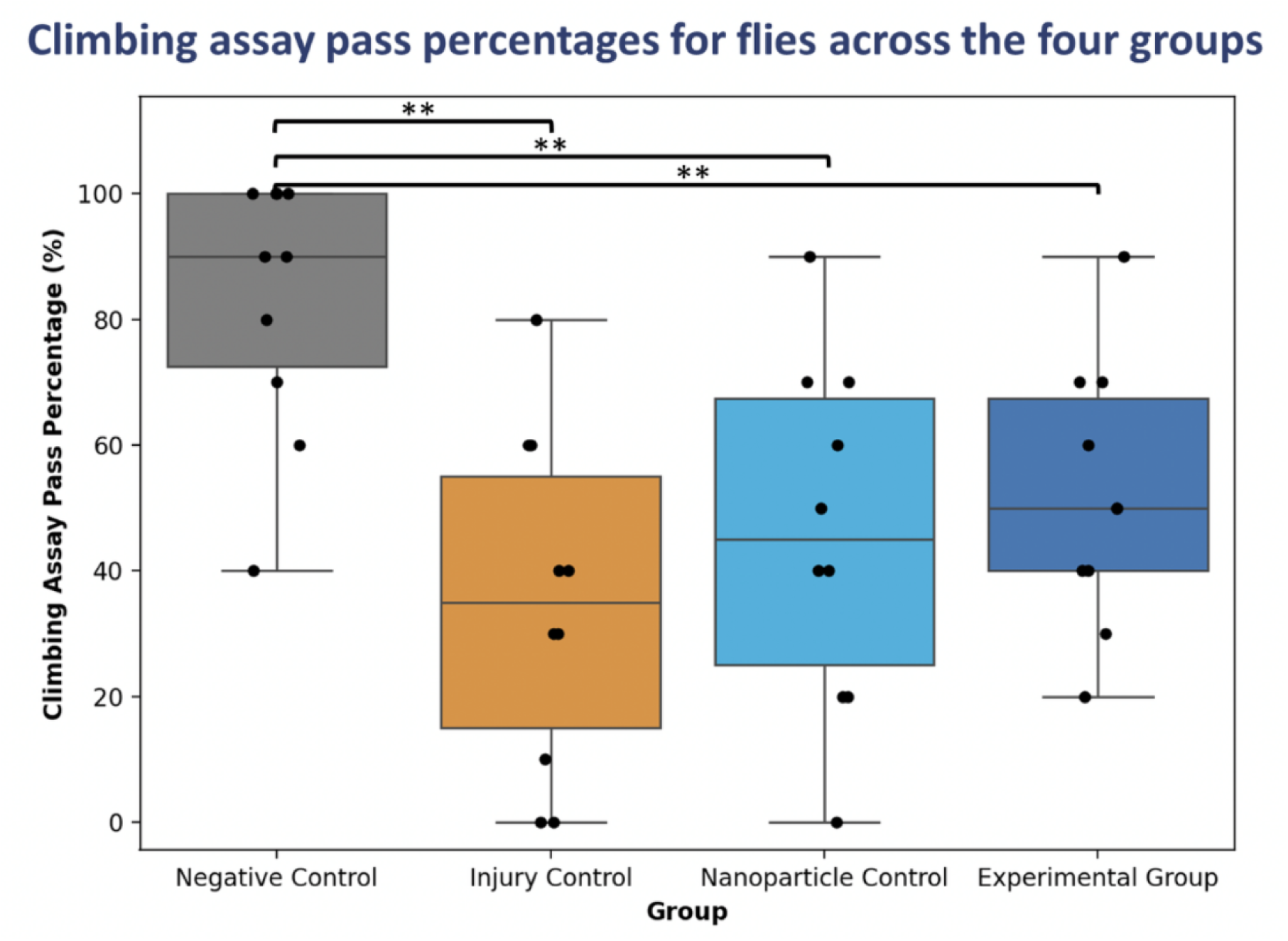
Negative geotaxis climbing assay results comparing motor function across the negative control, injury control, nanoparticle control, and experimental scaffold group. The injury control showed reduced climbing compared with the negative control, validating the TBI model, while the nanoparticle and scaffold groups showed non-significant increases in median pass rate compared with the injury control. ** indicates p<0.01 in a Mann-Whitney U comparison.

The median pass percentages were 90% for the negative control group, 35% for the injury control group, 45% for the nanoparticle control group, and 50% for the experimental scaffold group. Mann-Whitney U tests were used due to the small sample size being below 30 and thus not meeting the conditions of the Central Limit Theorem to use a t-distribution. Close medians and large ranges were present among groups 2, 3 and 4 leading to p-values above 0.05 in comparisons among the three groups. The procedure had a mortality rate of 45.2%. Individual trial pass rates ranged from 40% to 100% in the negative control, 0% to 80% in the injury control, 0% to 90% in the nanoparticle control, and 20% to 90% in the experimental scaffold group, indicating the substantial within-group variability present across all TBI groups.

## Discussion

The original purpose of the study was to investigate the effects of an electrospun piezoelectric hydrogel scaffold on motor function when applied to *Drosophila melanogaster* with traumatic brain injury. The purpose was addressed by fabricating the scaffold and applying it to flies, comparing motor function across groups through statistical analysis of climbing assay pass rates. Since the p-values of 0.0076, 0.0060, and 0.0014 are all less than the significance level of 0.05, the null hypothesis that the negative control group has the same median climbing assay pass rate as the groups afflicted with TBI was rejected. Thus, the different, higher, median climbing assay pass rate for the negative control group without TBI was shown to be statistically significantly different than the groups with TBI inflicted. This validated the protocol for inflicting traumatic brain injury, since the groups of flies with traumatic brain injury, particularly the injury control group that underwent TBI and a sham procedure, had statistically lower motor function, as expected according to the hypothesis and TBI protocol.

Since the p-values of comparisons between the injury control, nanoparticle control, and the experimental group, are all greater than the significance level of 0.05, the null hypothesis that the groups afflicted with TBI had the same median climbing assay pass rate was unable to be rejected. There was not a statistically significant difference between the groups found, and the results could have been due to random sampling variation. That being said, a clear trend emerged showing increased median climbing assay pass rates when flies had the scaffold applied (50%) compared to both when the fly had only nanoparticles applied (45%) and a sham procedure (35%).

There were many possible errors and assumptions made throughout the process of investigating the hypothesis. The main limitation of the study was the low sample size, with only 10 flies in every group. Typically, each vial of flies is used as one trial, while this study used each individual fly as one trial and calculated a pass percentage after multiple attempts in the climbing assay for the same fly. The low sample size was likely the root cause of the wild variance, along with the inconsistent application of traumatic brain injury. Although the varying severities of traumatic brain injury are more indicative of real world impact, not accounting for brain injury severity as a confounding variable and separating investigations on multiple severities, along with the low sample size that limited statistical findings, lead to a large variance in all of the groups. To ameliorate the issue, increasing the sample size and accounting for confounding variables by noting which flies had different severities of TBI before materials are applied could be promising future venues to reduce error.

Another error that limited the study was that the long term effects of the treatments and traumatic brain injury were not investigated. The justification for the hypothesis was long term regeneration and improved biocompatibility of the scaffold, but only 2-3 days were left between inflicting TBI and the procedure onto the flies and actually testing their motor function in a climbing assay. Analyzing results over a longer period of 2-3 weeks instead could have yielded different results, and the short timeframe might explain the marginal improvements from the scaffold that are not statistically significant but are part of a promising trend. Over time, the difference between the scaffold and the nanoparticle control and the injury control group could possibly be more pronounced, suggesting that more investigation is necessary.

Finally, one other assumption made was that survivorship bias would not affect the results. The high mortality rate from the procedure was due to the intensive experience of both traumatic brain injury and the procedure to open and seal the fly cuticle. The assumption that survivorship bias was insignificant was made due to the fact that all groups underwent a procedure to open and seal fly cuticles and were placed in a high impact trauma apparatus. Thus, only flies that could survive the intensive procedure were tested on, which could be less representative of other species like other animal models or humans that may not have such a high mortality rate due to not being as delicate as flies. The survivorship bias could have been improved by creating a more standardized method to open and close fly cuticles as well as using more sophisticated technology like robotic tools with more precision. Alternatively, another, more human-like model organism could have been used to more easily conduct the procedure.

Current traumatic brain injury treatments largely neglect fast and effective restorative and regenerative therapies, focusing on managing symptoms rather than systemic cell death and secondary changes that alter brain chemistry. While individually, piezoelectric materials, particularly nanoparticles, are emerging as a promising method to improve regeneration following traumatic brain injury in a way that can be remotely activated mechanically, the cytotoxic effects of some materials used in the field and the potential for agglomeration or a suboptimal microenvironment suggests that research is needed for a delivery mechanism that supports biocompatible and long term regeneration. Thus, the promising trend seen with an electrospun piezoelectric hydrogel scaffold found in the study is a potential new direction for safe and effective therapeutic restoration after a traumatic brain injury. The trend also suggests that combining the prominent methods in the field used in this study is possible and seems to not significantly hinder therapeutic effects, which is important for potential new avenues of maximizing therapeutic efficiency.

The trend found in the study suggests that future work should be conducted to investigate the merits of an electrospun piezoelectric hydrogel scaffold for traumatic brain injury more closely. Studies that use other model organisms such as animals without a hard covering over the brain like *C. elegans* or larger animals like rats that can be more easily cut into could be a promising future direction to achieve a larger sample size and less survivorship bias and evaluate the merits of the potential benefits of the scaffold. Furthermore, future research could focus on the mechanistic actions of the scaffold and the mechanical and electrical properties of it compared to just piezoelectric nanoparticles or other treatments. For example, studies with cell culture investigating the potential ROS scavenging or the activation of calcium-gated voltage ion channels that the scaffold can induce when stimulated by ultrasound could shed light on the reasons the scaffold should be expected to improve motor function.

Finally, future work can explore the merits of the electrospun piezoelectric hydrogel scaffold for use in other conditions and behavioral issues than just motor function after a traumatic brain injury. For example, traumatic brain injury leads to many cognitive deficits like memory loss along with the motor deficits, suggesting that evaluating memory or cognitive capabilities after a scaffold is applied could be a promising work. Furthermore, the similar cell death and need for a healing microenvironment is present in many neurodegenerative diseases like Alzheimer’s Disease, Parkinson’s Disease, Amyotrophic Lateral Sclerosis, as well as mental illness like depression and other acute experiences like stroke. Thus, the successfully synthesized and promising electrospun piezoelectric scaffold can be tested in other models or diseases for further evaluation of therapeutic potential.

The original hypothesis of the study was that the electrospun piezoelectric hydrogel scaffold would significantly improve motor function when applied to *Drosophila melanogaster* with traumatic brain injury as compared to the control groups that also had traumatic brain injury, including a group with only piezoelectric nanoparticles applied and a group with sham treatment applied. The successful synthesis of electrospun piezoelectric hydrogel scaffold with piezoelectric barium titanate nanoparticles and electrospun sodium alginate and polyethylene oxide formed a hydrogel when crosslinked with minimal beading. While the hypothesis was not statistically supported, the general trends that were found with increased median climbing assay pass rates for the experimental group and nanoparticle control group and particularly for the experimental group, supported the hypothesis. Thus, while the hypothesis cannot be supported, the promising trend suggests that future work that focuses on increased sample sizes and mechanistic research could be effective.

## Methods

### Variables

The independent variables were whether or not the flies received TBI or sham injury, in which the *Drosophila melanogaster* underwent the same HIT model as the TBI group but was not actually impacted, and whether they received either no scaffold, just ultrasound-stimulated piezoelectric nanoparticles, or an electrospun piezoelectric hydrogel scaffold stimulated by ultrasound. The dependent variable was the motor function of the *Drosophila melanogaster* as measured by the negative geotaxis climbing assay. Measuring motor function is particularly useful, since locomotor ability is one of the more consistently impaired behaviors after TBI in *Drosophila melanogaster*. An improvement in climbing performance was used to indicate a possible positive correlation between the scaffold and neural repair and functional recovery.

### Constants

The constants used in the experiment were the original fly state (Oregon-R wild-type *Drosophila melanogaster*), food fed to flies (mixture of corn syrup, yeast, soy flour, cornmeal, agar, and distilled water), lab equipment, time of day (between 1:45 PM and 3:30 PM EST), location (Dr. Eliason’s Room at the Academies of Loudoun - 2419), sex of tested flies (male), temperature of rearing (22°C for life cycle of 14 days), light cycle (12 hr light/12 hr dark cycle), the age at HIT (TBI/sham) (∼4 days post-eclosion for all flies) and the climbing assay (2-4 days after HIT for all flies), the TBI apparatus (HIT model) applied to all flies, and the procedure to open and reseal the cuticle of the flies.

### Controls

Five control groups were possible with the experiment: the negative control, two positive controls, and two toxicity controls. The negative control included wild-type *Drosophila melanogaster* that were put through the high impact trauma (HIT) apparatus without being actually harmed and that had their cuticles cut and glued with nothing applied to the brain, suggesting that the group should have performed well on the climbing assay due to the negative geotactic tendencies of *Drosophila melanogaster* (Dollinger et al., 2017). The negative control served as a baseline measurement for normal climbing and functions as a sham-surgery control, since it underwent the same cuticle opening/sealing steps without any materials applied, isolating the effect of procedural stress on climbing.

The first positive control included wild-type *Drosophila melanogaster* that had been afflicted with TBI through high-impact mechanical shock in the high-impact trauma (HIT) model, which could have cascaded into downstream secondary effects and impair motor skills. The use of this model involved evaluation of improvement through a climbing assay, in which performance should have been severely impaired (Katzenberger et al., 2013). The group should also have had their cuticles cut and glued with nothing applied. The first positive control was meant to be a group of flies with motor impairments poor performance on the climbing assay, serving as a model of TBI with no treatment. The second positive control included wild-type *Drosophila melanogaster* that had been hit in the HIT model, suggesting that there should have been motor impairment from TBI, and that they should have had their cuticles cut and glued with just BTNPs applied. The second positive control was meant to be a test of a previous treatment, with potentially slightly better motor function than the first positive control expected since a similar material had some positive effects in rats. The group served as a comparison to see if the potential effect can be improved in the experimental group.

The first toxicity control should have included wild-type *Drosophila melanogaster* that had been in the HIT apparatus without being hit, but that had their cuticles cut and glued with just BTNPs applied. The first toxicity control should have served as a method to test toxicity of the nanoparticles by seeing if motor function is impaired compared to what should be healthy flies in the negative control. Finally, the second toxicity control should have involved wild-type *Drosophila melanogaster* that were placed in the TBI apparatus without being hit, but that had their cuticles cut and glued with the full electrospun piezoelectric hydrogel scaffold applied. The second toxicity control should have been meant to investigate potential negative or unsafe impacts that the intended treatment may have by comparing motor function to what should be healthy flies in the negative control group. However, the toxicity controls did not relate very much to TBI itself, which means they were not very aligned with the purpose of the experiment and were deprioritized and eventually not tested. The three controls of the negative control and two positive controls were determined to be enough to come to some conclusions, so four total groups were used due to time constraints.

### Fly care and rearing

Oregon-R wild-type *Drosophila melanogaster* (Carolina Biological Supply #172100) were maintained at 22°C on a 12 hr light/12 hr dark cycle in clear plastic vials containing standard fly food (corn syrup, yeast, soy flour, cornmeal, agar, distilled water, and 10% propionic acid). Food was replaced when cracked or dry. Stocks were expanded by tapping flies into new vials every four days or maintained by tapping every three weeks. Male flies were isolated by cold sorting at 2-4 days post-eclosion for use in all experiments.

### Scaffold fabrication

Barium titanate nanoparticles (BTNPs; NanoAmor, tetragonal, ∼300 nm) were homogenized by sonication at 8W for 10 min and allowed to rest for 30 minutes. A 10 mL electrospinning solution was prepared by dispersing 20 mg (0.2 w/v) BTNPs in deionized water under sonication for 30 minutes, followed by dissolution of 120 mg sodium alginate (SA) and 180 mg polyethylene glycol (PEO) under continuous stirring for 24 hr. When BTNPs were visibly settled, the solution was loaded into a 10 mL syringe with a 20 G needle and electrospun at 15 kV, a flow rate of 0.3 mL/hr, and a 15 cm needle-to-collector distance. Collected nanofibers were dried at 50°C and then vacuum-dried for at least 3 days. The final scaffold was characterized by scanning electron microscopy (SEM) after sputter coating with gold and argon, and fiber morphology was analyzed using FiberMetric and Phenom software.

### TBI induction

TBI was induced using the high-impact trauma (HIT) apparatus (Katzenberger et al., 2013). The apparatus consisted of a wooden board with a spring mechanism fitted with a plastic vial and a polyurethane impact pad positioned beneath it. Approximately 50 unanesthetized flies were confined to the bottom quarter of a plastic vial with a cotton ball to restrict movement during impact. The vial was attached to the spring and deflected a 90° angle before being released to slam onto the polyurethane pad, delivering a high-impact shockwave through the flies to simulate closed-head TBI. This impact protocol has been shown to induce secondary injury cascades including oxidative stress and neuroinflammation, and to produce measurable motor deficits in the climbing assay (Katzenberger et al., 2013). The negative control group underwent the same handling and vial attachment procedure but was not exposed to any impact (sham injury), isolating the effect of TBI from procedural handling stress.

### Treatment application

Individual flies were anesthetized via CO₂ and secured in a 3D-printed fly sarcophagus (127 mm × 127 mm × 3 mm, with a 3 mm × 1 mm cutout). CO₂ anesthesia was used rather than cold temperatures, since cold temperatures at this scale would require individual surgical manipulation that was not feasible. However, CO₂ exposure was kept to a minimum in order to minimize potential neurological effects. Under a dissecting microscope, a triangular flap was cut in the head cuticle using microscissors, leaving a part of the cuticle intact to act as a hinge. Fine forceps were used to gently lift the flap and expose the underlying tissue. Depending on group assignment, the *Drosophila melanogaster* were exposed to approximately 20 µL of BTNP suspension (nanoparticle control), a 0.5 mm x 0.5 mm piece of electrospun scaffold (experimental group), or no material (negative and injury controls) on the exposed area. The scaffold piece was cut from the dried electrospun mat using fine scissors under the dissecting microscope to achieve the target dimensions. The *Drosophila melanogaster* were then exposed to a minimal amount of UV glue measured with a fine-tip applicator to reseal the cuticle flap. The glue was cured for 20 seconds under UV light at 1 cm distance. Flies were allowed to recover for 5 minutes. Survival and post-recovery locomotion were assessed, and any fly unable to stand or walk was excluded. A 45.2% procedural mortality rate was observed across all flies undergoing the cuticle-opening procedure, which substantially reduced the effective sample size in the treatment groups. For ultrasound activation, the fly in a sealed vial was exposed to 1 MHz ultrasound for 30 seconds using a Richmar US 1000 3rd Edition unit. Ultrasound was applied to both the nanoparticle control and experimental groups, as the piezoelectric mechanism requires mechanical stimulation to generate the electrical output hypothesized to drive neural recovery.

### Negative geotaxis climbing assay

Ten male flies per group were stored in vials at 22°C for 2-4 days following TBI induction and treatment before testing, allowing secondary injury cascades to develop (Katzenberger et al., 2013). The experiment was double-blinded by having an impartial classmate randomly assign vial numbers 1-4 and shuffle them prior to testing, so that the researcher scoring the assay did not know group assignments until after all data were collected. Individual flies were tapped into a 100 mL graduated cylinder marked at 5 cm above the base. The cylinder was tapped twice to bring the fly to the bottom, and the fly was given 30 seconds to cross the 5 cm mark. This was repeated 10 times per fly across 10 flies per group (N=10 trials per group). Each fly therefore contributed one pass rate value (number of passes out of 10 technical replicates) as the unit of analysis. The proportion of times each fly crossed the mark in 30 seconds was recorded as that fly’s pass rate. Flies that crossed and fell back below the line were still recorded as a pass. Flies on the line 30 seconds were not. This assay exploits the innate negative geotaxis behavior of *Drosophila melanogaster*, in which healthy flies reliably climb upward when tapped to the bottom of a vial. Impairment of this behavior following TBI reflects deficits in motor coordination and locomotor ability (Dollinger et al., 2017). Flies were euthanized in 70% ethanol at -20°C after testing.

### Statistical analysis

Pass rate data for each fly (proportion of 10 technical replicates resulting in a pass) were compiled across the 10 flies per group. Due to the small sample size and non-normal data distribution, pairwise Mann-Whitney U tests were used for group differences. The Mann-Whitney U test was selected because it does not assume normality and is appropriate for comparing medians across more than two independent groups with small sample sizes. A significance threshold of p < 0.05 was used for all comparisons. Because multiple pairwise comparisons were conducted, the probability of a Type 1 error across comparisons is elevated. Bonferroni or Holm corrections were not applied given the exploratory nature of this study, and this should be considered when interpreting the results.

### Overall procedure

After synthesizing the scaffold, each group was formed based on whether they had TBI or sham TBI and whether they were given a sham procedure, procedure with only barium titanate nanoparticles, or a procedure with the full scaffold.

**Figure 5.**
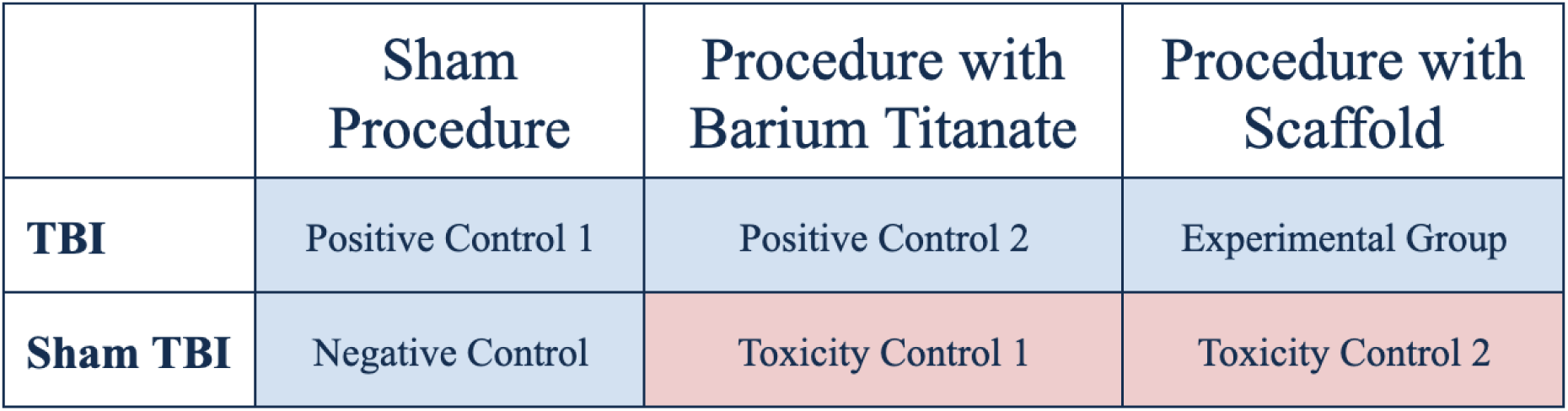
Overview of the four study groups used to evaluate motor recovery after TBI. The negative control underwent sham injury, the injury control received TBI without treatment, the nanoparticle control received TBI with ultrasound-stimulated BTNPs, and the experimental scaffold group received TBI with the ultrasound-stimulated SA/PEO/BTNP scaffold. The groups highlighted in red (toxicity controls) were not used due to time constraints.

After 2-3 days, the groups were then put under a negative geotaxis climbing assay to assess motor function, with the results recorded and statistically analyzed.

**Figure 6.**
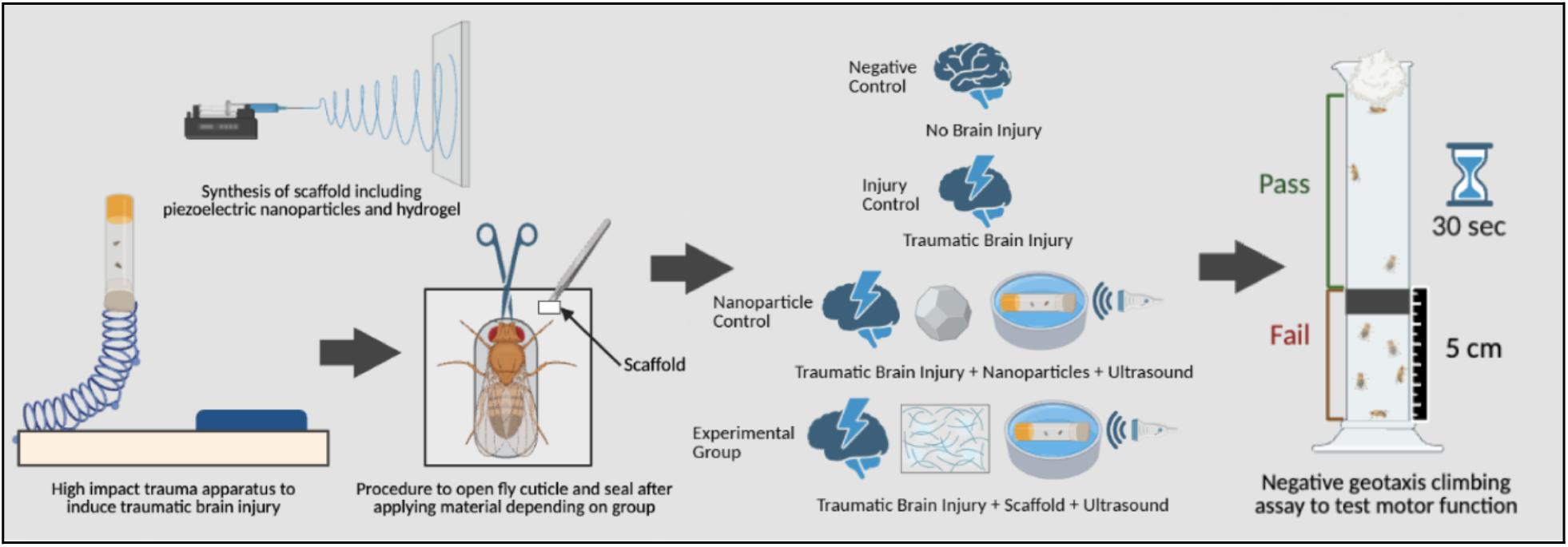
Schematic of the experimental workflow, including scaffold fabrication, fly rearing, TBI induction with the HIT apparatus, treatment application through cuticle opening, ultrasound stimulation, and negative geotaxis climbing assay testing after recovery. Created with BioRender.

### Safety

In accordance with the use of chemicals and flies, multiple safety protocols were followed. No eating or drinking in the lab was allowed and food and beverages were kept away from lab equipment, chemicals, or glassware to prevent contamination. Closed toed shoes were worn at all times in case of an accident. When working with chemicals and glassware, personal protective equipment (PPE) such as goggles and lab coats were worn at all times. Wearing hot hands during fly food preparation and maintaining the seal on the vials when tapping ensured that fly husbandry was safe and efficient. Both barium titanate nanoparticles and propionic acid can be toxic and irritating to the skin and eyes, which is why gloves and goggles were worn at all times when using them (NanoAmor, 2026; ThermoFisher Scientific, 2025a). Sodium alginate and polyethylene oxide were less hazardous, but still handled with care (ThermoFisher Scientific, 2023; ThermoFisher Scientific, 2025b). With all chemicals used, proper PPE was donned and hands were washed thoroughly before and after handling to prevent contamination. Furthermore, if chemicals were inhaled or ingested, a medical professional was contacted while giving researchers exposure to air. If chemicals contacted the skin or eyes, they were thoroughly washed with soap and water or an eye-wash station.

## Acknowledgements

Dr. Jessica Eliason, PhD mentored and guided the researchers through the project, including helping secure materials and providing feedback.

